# Discovery of a new *Phenuivirus* species in *Lachenalia* plants reveals possible co-evolution between 5’ and 3’ RNA sequence motifs

**DOI:** 10.1101/2024.09.04.611151

**Authors:** Rob J. Dekker, Wim C. de Leeuw, Marina van Olst, Wim A. Ensink, Selina van Leeuwen, Timo M. Breit, Martijs J. Jonker

## Abstract

This study reports the discovery of a new Phenuivirus species, named Lachenalia Phenuivirus-1 (LacPhV-1), from Lachenalia plants in an urban botanic garden in Amsterdam. Using a combination of smallRNA-seq, RNA-seq, and advanced bioinformatics, we identified a segmented negative-strand RNA virus in the Phenuiviridae family. Our findings show significant divergence between this new virus and known Phenuiviruses, such as Tulip Streak Virus (TuSV) and Lactuca Big Vein associated Phlebovirus (LBVaPV), supporting its classification as a distinct species. Notably, the sequence differences found in the conserved 5’ and 3’ ends of these segments suggest potential co-evolution. Despite the observed genomic distances, there is significant conservation in the RdRp subdomain, underscoring evolutionary relationships among LacPhV-1, TuSV, and LBVaPV. Our findings expand the known global virome and highlight the importance of exploring plant viromes in diverse ecological settings to better understand virus evolution and diversity.

## Introduction

The global virome is still largely unknown (Paez-Espino *et al*., 2016; Neri, *et al*. 2022). As a first step to understand the innumerable strategies viruses possess to evade the host anti-virus responses, it is important to discover as many different viruses as possible. This applies to human-pathogenic viruses (Carroll *et al*., 2018a and 2018b), as well as to phytopathogenic viruses, which are a serious threat to global food security (Jones, 2021; Ristaino *et al*., 2021). A particular environment that might be a breeding ground for, yet unknown, phytoviruses is the (urban) botanic garden. Given the green “islands” botanic gardens represent, in which plants are kept in relative seclusion for numerous generations, often in close proximity to exotic plant species from exotic places, the assumption is that there may be many (yet unknown) viruses present in botanic garden plants.

There are several ways to discover new phytoviruses. The most straightforward approach is to analyze plant RNA with RNA-seq and using bioinformatics to assemble the reads into contigs. The genome sequence of these contigs or their translated protein sequences can be compared to virus databases to detect known or yet unknown virus sequences. One problem with the RNA-seq approach is the massive background of host RNA. Also, past virus infections cannot be detected by lack of viral RNA. Another approach is to investigate the virus-related siRNA defense response of the host plants to a viral infection (Vaucheret *et al*., 2024). Given the massive siRNA response of most plants to a virus infection, which usually covers the entire length of RNA viruses, it is often possible to easily reconstruct the complete virus RNA sequence from the host-generated siRNAs (Wu *et al*., 2010). Moreover, the siRNA response can be present after the virus is defeated by the host, so siRNA might sometimes also provide information of past virus infections. A disadvantage of the siRNA approach relates to virus infections with multiple variants. As the typical length of siRNAs is 21 nucleotides, it is often difficult to entangle multiple virus variant sequences. This is obviously a lesser problem with paired-end 150 nucleotide RNA-seq reads. Hence, combining smallRNA-seq and RNA seq in virus discovery experiments will provide the best of both approaches.

To investigate the current or past presence of known virus variants or unknown viruses in botanic garden plants, we screened 25 *Asparagales* samples from a Dutch urban botanic garden by high-throughput smallRNA-sequencing, as well as RNA-sequencing. In this report we describe the discovery of a new segmented, negative-strand RNA *Bunayvirus* that appears to belong to the *Phenuiviridae* family. The sequence differences found in the 5’ and 3’ conserved ends of the four virus RNA segments hint at the possible presence of sequence co-evolution. This new virus was discovered along with two other newly identified viruses within the same sample set (Dekker *et al*., 2024a and 2024b).

## Material and methods

### Samples

Samples of leaves from 25 *Asparagales* plants were collected from Hortus Botanicus, a botanic garden in Amsterdam, the Netherlands, on February 14, 2019. Details about the plant genera can be found in Supplemental Table ST1.

### RNA isolation

Small-RNA was isolated by grinding a flash-frozen ±1 cm^2^ leaf fragment to fine powder using mortar and pestle, dissolving the powder in QIAzol Lysis Reagent (Qiagen) and purifying the RNA using the miRNeasy Mini Kit (Qiagen). Separation of the total RNA in a small (<200 nt) and large (>200 nt) fraction, including DNase treatment of the large RNA isolates, was performed as described in the manufacturer’s instructions. The concentration of the RNA was determined using a NanoDrop ND-2000 (Thermo Fisher Scientific) and RNA quality was assessed using the 2200 TapeStation System with Agilent RNA ScreenTapes (Agilent Technologies).

### RNA-sequencing

Barcoded smallRNA-seq and RNA-seq libraries were generated using a Small RNA-seq Library Prep Kit (Lexogen) and a TruSeq Stranded Total RNA with Ribo-Zero Plant kit (Illumina), respectively. The size distribution of the libraries with indexed adapters was assessed using a 2200 TapeStation System with Agilent D1000 ScreenTapes (Agilent Technologies). The smallRNA-seq libraries from samples S01 to S12 and from samples S14-S26 were clustered and sequenced at 2×75 bp and 1×75 bp, on a NextSeq 550 System using a NextSeq 500/550 Mid Output Kit v2.5 or a NextSeq 500/550 High Output Kit v2.5 (75 cycles or 150 cycles; Illumina), respectively. RNA-seq libraries were clustered and sequenced at 2×150 bp on a NovaSeq 6000 System using the NovaSeq 6000 S4 Reagent Kit v1.5 (300 cycles; Illumina).

### Bioinformatics analyses

Sequencing reads were trimmed using trimmomatic v0.39 (Bolger *et al*., 2014) [parameters: LEADING:3; TRAILING:3; SLIDINGWINDOW:4:15; MINLEN:19]. Mapping of the trimmed reads to the NCBI virus database was performed using Bowtie2 v2.4.1 (Langmead *et al*., 2012). Contigs were assembled from smallRNA-seq data using all trimmed reads as input for SPAdes De Novo Assembler (Prjibelski *et al*., 2020) with parameter settings: only-assembler mode, coverage depth cutoff 10, and kmer length 17, 18 and 21. Assembly of contigs from RNA-seq data was performed with default settings.

Scanning of contig sequences for potential RdRp-like proteins was performed using PalmScan (Babaian *et al*., 2022) and LucaProt (Hou *et al*., 2023).

## Results and discussion

### A new *Phenuivirus* in *Lachenalia*

In our study, we employed a combination of smallRNA-seq, RNA-seq, advanced bioinformatics, and manual curation to uncover previously unknown plant viruses in 25 *Asparagales* plants showing mild to severe disease phenotypes collected from an urban botanic garden in Amsterdam (Supplemental Table ST1). The smallRNA-seq experiment yielded on average approximately 11 million sequencing reads (< 76 nt) per sample and the RNA-seq experiment yielded on average about 39 million read-pairs (2×150 nt) per sample (Supplemental Table ST1).

Initial assembly of the smallRNA-seq reads resulted in several contigs that showed a weak similarity to the RdRp containing RNA1 segment of Tulip Streak Virus (TuSV) (Neriya *et al*., 2021). TuSV is a four-part segmented negative-stranded, single-stranded RNA Phenuivirus from the order *Bunyavirales* that initially was reported to contain two segments of ∼6 kb and 1.1 kb (Neriya *et al*., 2021), but was by later research extended to include two additional smaller segments of ∼1.1 kb and 1.3 kb (Supplemental Table ST2). The TuSV-like siRNA-derived contigs, combined with the RNA-seq contigs were assembled into four virus RNA segments (RNA1-RNA4; GenBank accessions: PQ067367, PQ067368, PQ067369, PQ067370) that code for four virus-related proteins (Figure 1A and Supplementary Table ST2).

**Figure 1.**
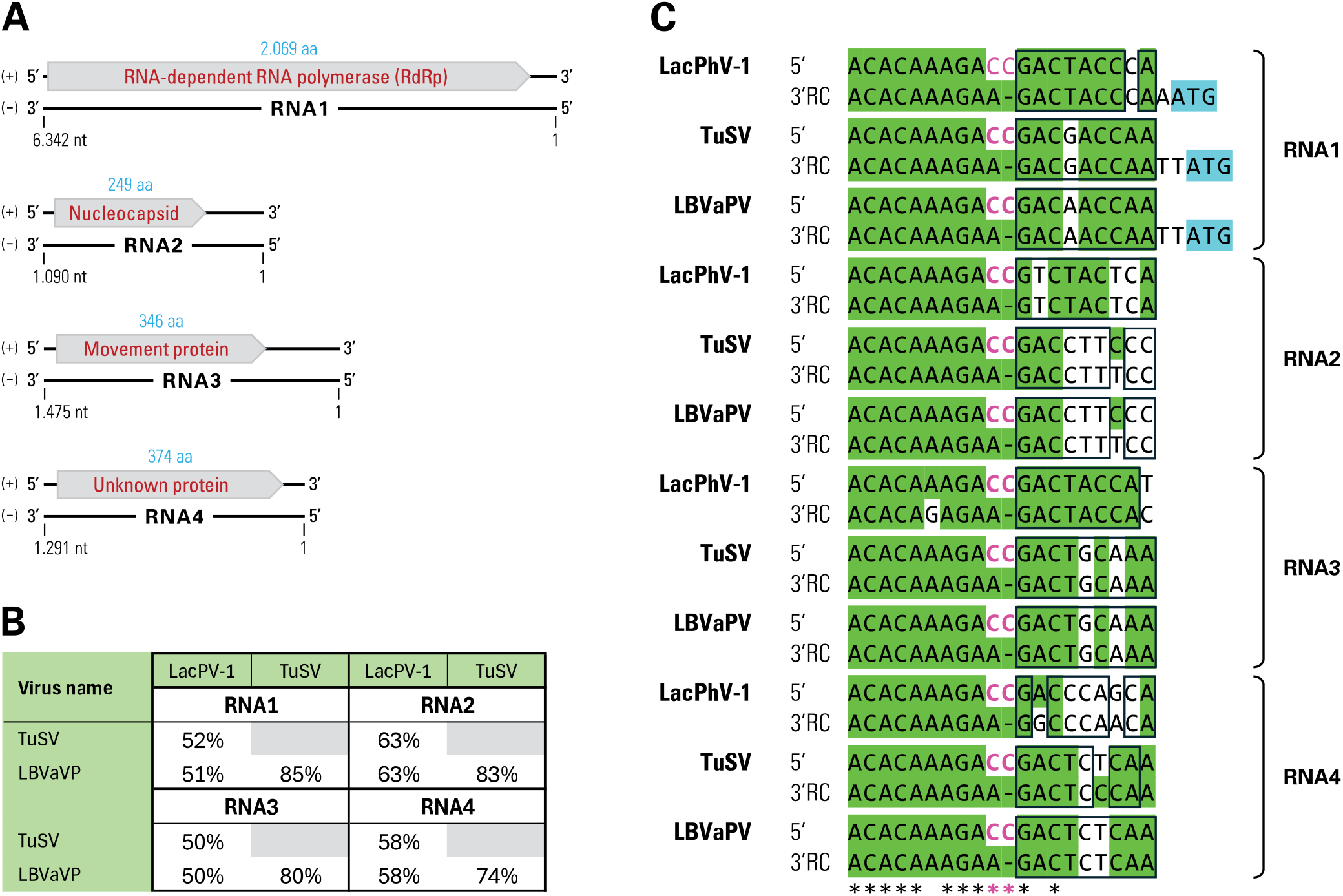
A) Schematic illustration of the discovered Lachenalia Phenuivirus-1 (LacPhV-1) genome and its associated proteins. Indicated on the + strand are the proposed open reading frames (grey arrow box) and corresponding protein sizes (blue). B) Amino acid similarity between (LacPhV-1-NL1-26) and the two most similar Phenuivirus genome sequences: Tulip Streak Virus (TuSV) and Lactuca Big Vein associated Phlebovirus (LBVaPV). C) Similarities between the 5’ and 3’ terminus motifs of the four genomic RNA segments of each virus. To facilitate interpretation of the sequence complementarity, the reverse complement of the 3’ terminal sequences are shown (3’RC). Nucleotides identical to the consensus nucleotide are highlighted in green. The hallmark non-complementary, protruding CC at positions 10 and 11 in the 5’ motif are indicated in pink (Ren *et al*., 2020). Nucleotides from positions 11 (3’) and 12 (5’) that are identical in the same RNA molecule are indicated by a black box. The ATG translation start sites for the RdRP proteins are highlighted in blue.

The similarity between the specific RNAs of this new Phenuivirus and the known TuSV virus ranges from 56-68% (at a coverage of 36-72%) at the RNA level and 50-63% (at a coverage 87-100%) at the protein level (Figure 1B and Supplemental Table ST2). These similarities are well below the species demarcation criteria for the *Phenuiviridae* family identity (< 95% identity in the RdRp amino acd sequence, Sasaya *et al*., 2023) to conclude that the virus sequences we found are from a yet unknown Phenuivirus species. When we mapped the smallRNA-seq reads and RNA-seq reads from all samples back to the new virus RNA sequences, we saw evidence for siRNA related to this virus in four samples (S09, S15, S22, S26), all of which were *Lachenalia* samples (Supplemental Tables 1 and 3). Hence, we named our new virus Lachenalia Phenuivirus-1 (LacPhV-1) and the isolation variant sequence from sample 26 was used in our subsequent analyses, LacPhV-1-NL1-26.

There was one other Phenuivirus in GenBank that also showed similarity to both LacPhV-1 and TuSV; Lactuca Big Vein associated Phlebovirus (LBVaPV) (Schravesande *et al*., 2023; Supplemental Table ST2). Even though TuSV and LBVaPV differ substantially, with a similarity of 69-75% at RNA level and 74-85% at protein level, they show almost the same weak similarity to LacPhV-1, which results in a triangular sequence distance from each other (Figure 1B and Supplemental Table ST2). This is clear at the protein level, where the similarity coverage is always at least 87%, whereas at the RNA level this coverage goes even as low as 22%. The fact that the overall RNA similarity is lower than protein similarity is a common occurrence in viral genome comparisons. Given that these three distinct viruses were found in plants from three different orders: Asparagales (LacPhV-1), Liliales (TuSV), and Asterales (LBVaPV), might explain the observed genomic distances.

When we zoomed in on the RdRp subdomain region with the well-conserved motif A, B, and C, which play an essential role in the catalytic function of RdRp (Jia and Gong, 2019), it was clear that while there was substantial difference between LacPhV-1 and TuSV (19%) or LBVaPV (19%), the three sequences of the three domains were identical (Figure 2) even though they represent 23% of the subdomain sequences. The next most similar RdRp subdomain (Arenavirus from the *Arenaviridae* family), contained divergent amino acids in each of the three domains (Figure 2), suggesting that although the LacPhV-1, TuSV and LBVaPV are different species, they are evidently related.

**Figure 2.**
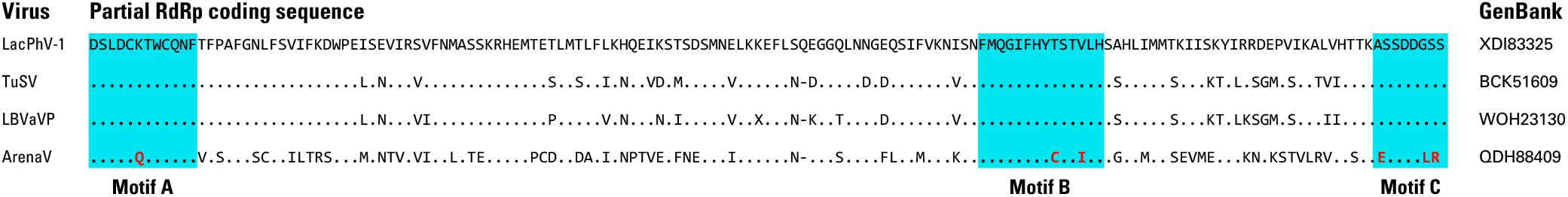
Sequence comparison of the conserved RdRp palm subdomains with the characteristic motifs. Alignment of the LacPhV conserved RdRp palm subdomain protein sequences to the three most similar RdRp subdomain sequences in GenBank. The amino acid sequences of the RdRp subdomain motifs A, B, and C are indicated in blue. Amino acids that diverge from the LacPhV-1 RdRp subdomain motifs are indicated in red.

We also observed a noticeable difference in relative smallRNA-seq read counts between the RNA segments of this new virus. After correction for read length, the relative read counts for the RNA1 and RNA 4 segments were 0.11 and 0.34 times the overall average read count, whereas this was 2.31 and 1.24 for the RNA2 and RNA3 segments, respectively (Supplemental Table ST4). This difference in smallRNA-read counts was consistent over all four LacPhV-1 positive samples. As most of those reads are 21-mers, this means that for unknown reasons the relative number of siRNAs for each RNA fragment is rather dissimilar. This phenomenon is also present with RNA-seq reads with low relative read counts (RNA1, 0.22 and RNA3, 0.78) and high relative read counts (RNA2, 1.91 and RNA4, 2.64). RNA-seq reads represent either virus RNA or virus mRNA. Hence, there is no direct correlation detectable between the relative number of siRNAs and virus (m)RNA (Supplemental Table ST4).

The four RNA segments of the LacPHV-1 virus contain complementary 5’ and 3’ terminal sequences that can form a so-called ‘panhandle structure’ which is typical for *Bunyaviruses* (Takahashi *et al*., 1990, Reguera *et al*., 2013 Sasaya *et al*., 2023). When we compare these terminal sequences with those of TuSV and LBVaPV, they show a stretch of nine nucleotides that is in all except one terminal sequence completely conserved (Figure 1D). The 5’ sequence continues with a conserved CC, whereas the 3’ sequence shows one T, leading to an important protruding C nucleotide when both ends pair (Ren *et al*., 2020). Comparing the sequence termini of these Phenuiviruses with other four or eight segmented Phenuiviruses (Supplemental Figure SF1) revealed, next to the hallmark non-complementary CC nucleotides, at nucleotide position 10 and 11 in the 5’ terminus motif, possible importance for positions 9 (5’ and 3’) and 10 (3’). These nucleotides invariably are (for the 3’ sequence in reverse complement orientation) ‘TC’ in Tenuiviruses and Mechloroviruses, ‘AC’ in Wenriviruses (Supplemental Figure SF1), and AA in the three somewhat similar Phenuiviruses in this study, of which the previously reported viruses are linked to Phleboviruses, even though this genus reportedly only has three segments (Sasaya *et al*., 2023).

In LacPhV-1, TuSV and LBVaPV, the second stretch of nine nucleotides in the terminal sequences seems to be more virus/RNA-segment specific, as no terminal sequence occurs in more than one RNA segment for all three viruses. Moreover, compared to the consensus sequence of this second nucleotide stretch, there are 33 non-consensus nucleotide pairs (31%) of the total 108 pairs. Yet more striking, only 6 nucleotide pairs (5.6%) are not complementary. Thus, even though there are quite some differences in the second stretch, there are few non-complementary divergences, which might be extremely important for vital virus functions as has been shown by Kohl *et al*. (2023) through the effects of a single point mutation in a 3’ terminus of a *Bunyavirus*. The inclination towards maintaining complementarity might indicate a form of co-evolution between the 5’ and 3’ termini of the RNA segment of these *Phenuiviruses*.

## Concluding remarks

With this study, we add a new virus species to the ever-expanding global virome. In our research into virus presence in (urban) botanic gardens, this is the second new virus species we report (Dekker *et al*., 2024). The first new virus species concerned a single-stranded, positive-stranded RNA virus of the *Capillovirus* genus and *Betaflexiviridae* family. Here we describe a single-stranded, negative-stranded RNA virus of the *Phenuiviridae* family.

The observation that the siRNA response to the four virus RNA segments, as well as the presence of these virus RNA molecules are both different and not related, yet quite consistent between samples is puzzling. Constant variation in the presence of virus RNA or virus mRNA is explicable; however, constant variation in virus-related siRNA hints at an siRNA regulatory mechanism based on the sequence of the RNA segments. Most likely, this is a cumulative effect of the siRNA hot-spots and cold spots in the various RNA segments (Molnár A *et al*., 2005).

With our new Phenuivirus we detected an indication of a possible presence of sequence co-evolution of the 5’ and 3’ termini that can form so-called panhandle RNA structures in the four molecules, probably involved in RNA replication, transcription, pseudo-circularization, and virus packaging (Barr *et al*., 2003 and 2004, Reguera *et al*., 2013,Sun *et al*., 2018, Ren *et al*., 2020, and Malet *et al*., 2023). Each addition of well-analyzed and well-annotated virus sequences to the virus databases increases the potential for new discoveries in the mechanisms of viruses, which may be beneficial to the battles against pathogenic viruses.

## Acknowledgements

We would like to express our sincere gratitude to Sarina Veldman, Martin Smit, Iris van Kleinwee and Reinout Havinga from the Hortus Botanicus in Amsterdam, The Netherlands, for their invaluable support in providing us with plant leaf material for this study. This research was directly and indirectly funded by the Swammerdam Institute for Life Sciences of the University of Amsterdam.

## Data availability

The raw sequence reads have been deposited in the NCBI Sequence Read Archive under BioProject accession number PRJNA1137160. The following Lachenalia Phenuivirus-1 (LacPhV-1) genome segment sequences have been deposited in NCBI GenBank: RNA1 (PQ067367), RNA2 (PQ067368), RNA3 (PQ067369), and RNA4 (PQ067370).

## Supplemental information

**Supplemental Figure SF1.**
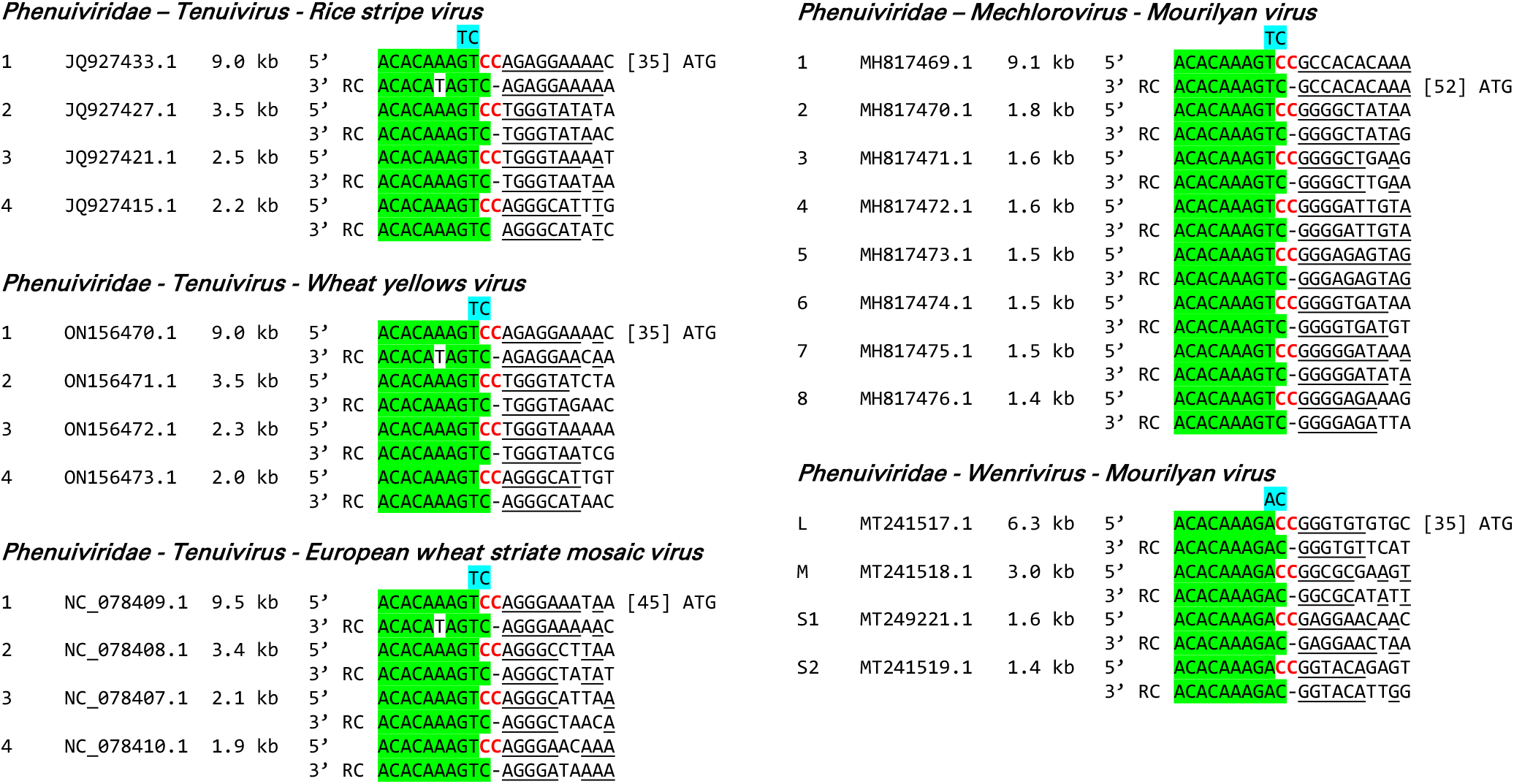
Comparison of Phenuiviridae segment termini. Similarities between the 5’ and 3’ terminus motifs of the four RNAs of each virus. To facilitate interpretation of the sequence complementarity, the reverse complement (RC) of the 3’ terminal sequences are presented. The consensus nucleotides from position 1 to 9 are highlighted in green. The hallmark CC nucleotides at positions 10 and 11 in the 5’ motif are indicated in red. The distinguishing nucleotides at position 9 (5’ and 3’) and position 10 (3’) are highlighted in blue. Nucleotides from positions 11 (3’) and 12 (5’) that are identical in the same RNA molecule are boxed.

**Supplemental Table ST1.** Overview of plant samples used for smallRNA and RNA-seq analyses. The table includes detailed information on the disease phenotype, harvest date, and taxonomic classification of each sample. Additionally, the number of reads obtained from smallRNA-seq and the number of read-pairs from RNA-seq are provided. (*) SmallRNA-seq was performed in two separate experiments, resulting in the observed variations in read counts.

| Sample ID | Name | Abbreviation | Pot number | Disease phenotype | Harvest date | Taxonomy |  |  |  | # of reads/read-pairs |  |
| --- | --- | --- | --- | --- | --- | --- | --- | --- | --- | --- | --- |
|  |  |  |  |  |  | Order | Family | Genus | Species | sRNAseq* | RNAseq |
| S01 | Drimiopsis maculata Lindl Paxton | Dm | 2 | moderate | 2019-02-14 | Asparagales | Asparagaceae | Drimiopsis | D. maculata | 3,570,212 | 34,640,313 |
| S02 | Lachenalia aloides Asch. Graebert var. quadricolor | La | 3 | severe | 2019-02-14 | Asparagales | Asparagaceae | Lachenalia | L. aloides | 5,955,615 | 38,542,291 |
| S03 | Lachenalia bulbifera Cirillo Engl | Lb | 4 | severe | 2019-02-14 | Asparagales | Asparagaceae | Lachenalia | L. Bulbifera | 4,975,216 | 54,405,719 |
| S04 | Lachenalia bulbifera Cirillo Engl | Lb | 13 | absent | 2019-02-14 | Asparagales | Asparagaceae | Lachenalia | L. Bulbifera | 4,498,177 | 48,864,675 |
| S05 | Albuca canadensis L. FMLeight | Ac | 6 | moderate | 2019-02-14 | Asparagales | Asparagaceae | Albuca | A. canadensis | 5,094,352 | 74,387,234 |
| S06 | Ferraria crispa Burm | FC | 7 | severe | 2019-02-14 | Asparagales | Iridaceae | Ferraria | F. crispa | 4,511,770 | 39,227,726 |
| S07 | Ferraria crispa Burm subsp. crispa | FC<sup>10</sup> | 10 | severe | 2019-02-14 | Asparagales | Iridaceae | Ferraria | F. crispa | 3,650,466 | 16,393,962 |
| S08 | Ferraria crispa Burm subsp. crispa | FC<sup>11</sup> | 11 | severe | 2019-02-14 | Asparagales | Iridaceae | Ferraria | F. crispa | 5,461,449 | 50,399,202 |
| S09 | Lachenalia liliflora Jacq | Ll | 14 | severe | 2019-02-14 | Asparagales | Asparagaceae | Lachenalia | L. liliflora | 5,193,916 | 41,434,192 |
| S10 | Albuca canadensis | Ac | 16 | moderate | 2019-02-14 | Asparagales | Asparagaceae | Albuca | A. canadensis | 4,000,657 | 31,769,287 |
| S11 | Tritonia pallida Ker. Gawl. subsp. pallida | Tp | 21 | moderate | 2019-02-14 | Asparagales | Iridaceae | Tritonia | T. pallida | 5,631,974 | 20,052,549 |
| S12 | Gladiolus splendens Sweet Herb | Gs | 22 | severe | 2019-02-14 | Asparagales | Iridaceae | Gladiolus | G. splendens | 5,061,686 | 36,381,503 |
| S14 | Drimiopsis maculata Lindl Paxton | Dm | 1 | absent | 2019-02-14 | Asparagales | Asparagaceae | Drimiopsis | D. maculata | 10,854,121 | 56,652,319 |
| S15 | Lachenalia aloides Asch. Graebert var. luteola | La | 5 | severe | 2019-02-14 | Asparagales | Asparagaceae | Lachenalia | L. aloides | 15,559,011 | 45,746,374 |
| S16 | Ferraria crispa Burm | FC | 7 | severe | 2019-02-14 | Asparagales | Iridaceae | Ferraria | F. crispa | 34,660,163 | 31,620,407 |
| S17 | Ferraria crispa Burm | FC | 8 | moderate | 2019-02-14 | Asparagales | Iridaceae | Ferraria | F. crispa | 21,189,604 | 22,966,473 |
| S18 | Ferraria crispa Burm | FC | 9 | severe | 2019-02-14 | Asparagales | Iridaceae | Ferraria | F. crispa | 11,280,321 | 32,402,472 |
| S19 | Ferraria crispa Burm subsp. crispa | FC<sup>10</sup> | 10 | severe | 2019-02-14 | Asparagales | Iridaceae | Ferraria | F. crispa | 23,454,445 | 21,800,891 |
| S20 | Ferraria crispa Burm subsp. crispa | FC<sup>11</sup> | 11 | severe | 2019-02-14 | Asparagales | Iridaceae | Ferraria | F. crispa | 15,185,230 | 69,625,722 |
| S21 | Ferraria crispa Burm subsp. crispa | FC | 12 | severe | 2019-02-14 | Asparagales | Iridaceae | Ferraria | F. crispa | 13,570,849 | 21,285,878 |
| S22 | Lachenalia stayneri WFBarker | Ls | 15 | moderate | 2019-02-14 | Asparagales | Asparagaceae | Lachenalia | L. stayneri | 15,290,142 | 37,664,978 |
| S23 | Albuca canadensis | Ac | 17 | moderate | 2019-02-14 | Asparagales | Asparagaceae | Albuca | A. canadensis | 11,732,233 | 24,542,074 |
| S24 | Albuca canadensis | Ac | 18 | moderate | 2019-02-14 | Asparagales | Asparagaceae | Albuca | A. canadensis | 10,911,507 | 49,361,870 |
| S25 | Lachenalia corymbosa JCManning Goldblatt | Lc | 19 | severe | 2019-02-14 | Asparagales | Asparagaceae | Lachenalia | L. corymbosa | 12,504,425 | 35,099,651 |
| S26 | Lachenalia kliprandensis WF. Barker | Lk | 20 | moderate | 2019-02-14 | Asparagales | Asparagaceae | Lachenalia | L. kliprandensis | 19,037,058 | 35,044,220 |
| Average number of reads |  |  |  |  |  |  |  |  |  | 10,913,384 | 38,812,479 |

**Supplemental Table ST2.** Sequence similarity comparisons between LacPhV-NL1-26, identified in this study, and various related viruses retrieved from GenBank. The table presents sequence similarity at both the genome and protein levels, with specific GenBank accession IDs provided for each virus and corresponding RNA genome segment. Pairwise comparisons are shown for LacPhV-NL1-26 against TuSV and LBVaPV, as well as a direct comparison between TuSV and LBVaPV (column TuSV vs LBVaPV).

| GENOME |  |  |  |  |  |  |  |  |  |  |  |  |
| --- | --- | --- | --- | --- | --- | --- | --- | --- | --- | --- | --- | --- |
| Segment | LacPhV-1-NL1-26 |  | TuSV |  |  |  | LBVaPV |  |  |  | TuSV vs LBVaPV |  |
|  | Accession | Size | Accession | Size | Coverage | Identity | Accession | Size | Coverage | Identity | Coverage | Identity |
| RNA1 | PQ067367 | 6,342 | LC571987.1 | 6,379 | 72% | 68.20% | OR610326.1 | 6,408 | 59% | 70.93% | 99% | 75.25% |
| RNA2 | PQ067368 | 1,090 | LC571988.1 | 1,110 | 46% | 67.01% | OR610327.1 | 1,107 | 42% | 69.74% | 100% | 74.98% |
| RNA3 | PQ067369 | 1,475 | LC805881.1 | 1,454 | 36% | 66.42% | OR610328.1 | 1,429 | 22% | 65.06% | 80% | 74.82% |
| RNA4 | PQ067370 | 1,291 | LC805882.1 | 1,283 | 60% | 65.26% | OR610329.1 | 1,282 | 67% | 65.63% | 100% | 69.26% |

| PROTEIN |  |  |  |  |  |  |  |  |  |  |  |  |
| --- | --- | --- | --- | --- | --- | --- | --- | --- | --- | --- | --- | --- |
| Segment | LacPhV-1-NL1-26 |  | TuSV |  |  |  | LBVaPV |  |  |  | TuSV vs LBVaPV |  |
|  | Accession | Size | Accession | Size | Coverage | Identity | Accession | Size | Coverage | Identity | Coverage | Identity |
| RNA1 | XDI83325 | 2,069 | BCK51609.1 | 2,095 | 99% | 57.47% | WOH23130.1 | 2,098 | 99% | 57.24% | 100% | 84.56% |
| RNA2 | XDI83326 | 249 | BCK51610.1 | 259 | 87% | 63.47% | WOH23131.1 | 260 | 88% | 62.73% | 100% | 83.40% |
| RNA3 | XDI83327 | 346 | BFI02998.1 | 348 | 100% | 49.86% | WOH23132.1 | 348 | 100% | 50.00% | 100% | 80.00% |
| RNA4 | XDI83328 | 374 | BFI02999.1 | 374 | 100% | 57.75% | WOH23133.1 | 374 | 100% | 58.02% | 100% | 73.80% |

**Supplemental Table ST4.** Relative read counts of smallRNA-seq and RNA-seq read mapping to the new LacPhV-1-NL1-26 RNA sequences. The table presents the number of reads mapping to each segment, normalized to segment length (per 1,000 nucleotides). Additionally, the normalized read counts are expressed as a ratio over the average read count per segment, providing a comparative analysis of the read distribution across the viral genome.

|  | Genome segment | Length (nt) | Read count per 1.000 nt |  |  |  | Relative to average segment read count |  |  |  |  |
| --- | --- | --- | --- | --- | --- | --- | --- | --- | --- | --- | --- |
|  |  |  | S09 | S15 | S22 | S26 | S09 | S15 | S22 | S26 | Average |
| smallRNA-seq | RNA1 | 6,342 | 2,636 | 25,295 | 23,048 | 21,841 | 0.05 | 0.11 | 0.10 | 0.20 | 0.11 |
|  | RNA2 | 1,090 | 116,469 | 548,200 | 568,169 | 234,458 | 2.33 | 2.35 | 2.41 | 2.16 | 2.31 |
|  | RNA3 | 1,475 | 66,308 | 281,544 | 283,555 | 132,263 | 1.33 | 1.20 | 1.20 | 1.22 | 1.24 |
|  | RNA4 | 1,291 | 14,347 | 79,661 | 69,230 | 45,782 | 0.29 | 0.34 | 0.29 | 0.42 | 0.34 |
|  | Average per 1.000 nt |  | 49,940 | 233,675 | 236,000 | 108,586 |  |  |  |  |  |
| RNA-seq | RNA1 | 6,342 | 22,441 | 7,063 | 40,933 | 25,442 | 0.45 | 0.03 | 0.17 | 0.23 | 0.22 |
|  | RNA2 | 1,090 | 178,862 | 68,673 | 501,528 | 177,967 | 3.58 | 0.29 | 2.13 | 1.64 | 1.91 |
|  | RNA3 | 1,475 | 67,552 | 19,563 | 176,409 | 79,193 | 1.35 | 0.08 | 0.75 | 0.73 | 0.73 |
|  | RNA4 | 1,291 | 221,149 | 78,137 | 814,953 | 254,821 | 4.43 | 0.33 | 3.45 | 2.35 | 2.64 |
|  | Average per 1.000 nt |  | 122,501 | 43,359 | 383,456 | 134,356 |  |  |  |  |  |

**Supplemental Table ST3.**
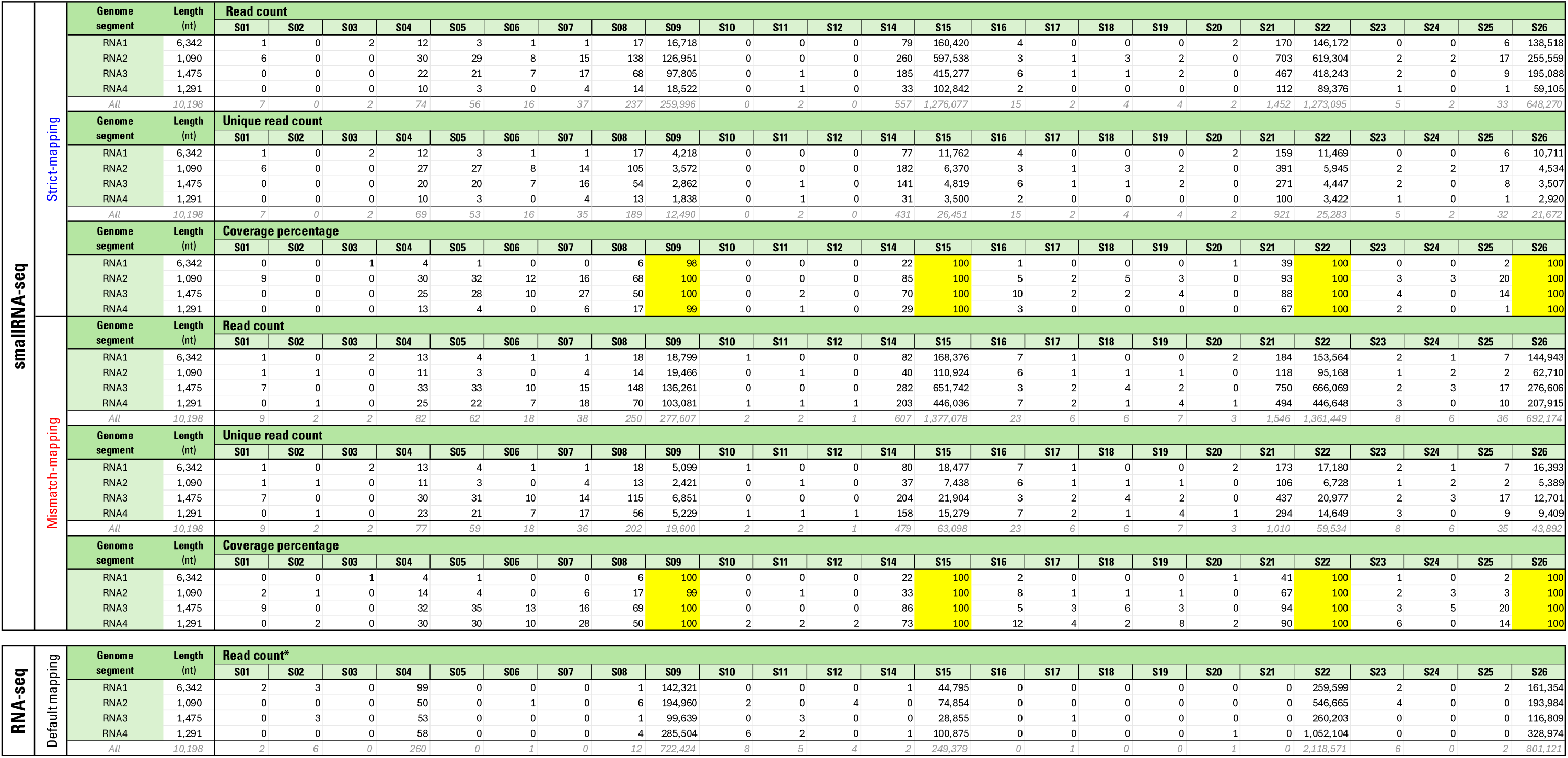
Summary smallRNA-seq and RNA-seq data mapping on the four genome segments of the discovered LacPhV-NL1-26 virus. SmallRNA-seq data was mapped using two different approaches: strict-mapping (allowing no mismatches; blue) and mismatch-mapping (allowing up to 10% mismatches; red). RNA-seq data was mapped using default settings. For each genomic segment (RNA1-4) of the LacPhV-NL1-26 virus, the table shows the read count, unique read count, and coverage percentage of the mapping. The “All” category aggregates the total number of reads mapped across all genomic segments. * The coverage percentage of the RNA-seq mapping is 100% for all four segments for all virus-positive samples.

